# Extracellular filaments revealed by affinity capture cryo-electron tomography

**DOI:** 10.1101/2023.08.05.552110

**Authors:** Leeya Engel, Magda Zaoralová, Momei Zhou, Alexander R. Dunn, Stefan L. Oliver

## Abstract

Cryogenic-electron tomography (cryo-ET) has provided an unprecedented glimpse into the nanoscale architecture of cells by combining cryogenic preservation of biological structures with electron tomography. Micropatterning of extracellular matrix proteins is increasingly used as a method to prepare adherent cell types for cryo-ET as it promotes optimal positioning of cells and subcellular regions of interest for vitrification, cryo-focused ion beam (cryo-FIB) milling, and data acquisition. Here we demonstrate a micropatterning workflow for capturing minimally adherent cell types, human T-cells and Jurkat cells, for cryo-FIB and cryo-ET. Our affinity capture system facilitated the nanoscale imaging of Jurkat cells, revealing extracellular filamentous structures. It improved workflow efficiency by consistently producing grids with a sufficient number of well positioned cells for an entire cryo-FIB session. Affinity capture can be extended to facilitate high resolution imaging of other adherent and non-adherent cell types with cryo-ET.

## MAIN

Cryogenic-electron tomography (cryo-ET) is a state-of-the-art electron microscopy (EM) modality used for the structural analysis of intact, vitrified cells at the nanometer and subnanometer scales ^1–6^. In preparation for cryo-ET, cells are deposited or grown on specialized substrates called EM grids. To image intracellular structures of eukaryotic cells, which are on the length scale of micrometers, thinning the cells using techniques such as cryo-focused ion beam (cryo-FIB) milling is typically required. Cells often distribute in a non-uniform manner on EM grids, and settle on the metal grid bars where they are not accessible for cryo-FIB and cryo-ET. In addition, cells can form clumps that prevent proper vitrification and successful FIB milling. These limitations present an obstacle both to the collection of cryo-ET data of sufficient quality from biological specimens and to the automation of data collection and cryo-FIB milling.

To gather sufficient data for high resolution structural determination, reproducible samples are needed. Time on high-end cryo-FIB and cryo-ET instrumentation, typically accessed through shared cryo- EM facilities, is expensive and limited. An ideal sample for cryo-FIB/cryo-ET would have a repeatable and uniform distribution of cells situated in the centers of the grid squares. We and others have published studies demonstrating the positioning of individual adherent cells or adherent cell pairs on EM grids micropatterned with extracellular matrix (ECM) proteins in preparation for cryo-ET ^7–12^. Another approach for providing a uniform distribution of cells for cryo-FIB/cryo-ET, applicable to both adherent and non-adherent cell types, is the Waffle Method ^13^. It employs high pressure freezing to vitrify a contained, continuous suspension of cells in preparation for cryo-FIB/cryo-ET. High pressure freezing, however, can be technically challenging and results in thicker samples (10s of microns) that require extended cryo-FIB milling times.

Lymphocytes (e.g., T-cells and B-cells) are minimally adherent cells that play important roles in the body’s defense against infectious diseases and cancer. Recent advances in our understanding of lymphocyte function have been provided by fluorescence imaging and conventional EM ^14,15^. However, the resolution of fluorescence imaging is limited by the wavelength of light and conventional EM techniques require fixation and staining steps that can distort or destroy delicate biological structures.

Here we present an affinity capture system that facilitates high-throughput cryo-FIB milling and subsequent nanoscale imaging by cryo-ET of non-adherent cell types that is compatible with vitrification by plunge-freezing. We micropatterned antibodies (Abs) onto EM grids to capture T-cells as well as Jurkat cells, an immortalized line of human T lymphocytes, and position them in the centers of EM grid squares. This facilitated our observation of nanoscale filaments emanating from the Jurkat cells. Subtomogram averaging (STA), RNA sequencing (RNA-seq), and flow cytometry suggest that these structures are intermediate filaments composed of vimentin. To our knowledge, this is the first time that micropatterning has been used to prepare minimally adherent cell types for cryo-ET.

## RESULTS

To assess the feasibility of capturing non-adherent cell lines on EM grids, a T-cell specific antibody to human CD3 (epsilon chain) was micropatterned to trap the T-cell surrogate, Jurkat cells, on islands within grid squares. Affinity capture grids were able to successfully capture single Jurkat cells and position them in the centers of the grid squares (**Fig. 1A to 1C, S1, and S2**). Cell localization was restricted to the micropatterned region on EM grids treated with the anti-CD3 antibody (**Fig. 1B and 1C**). Neither of the control conditions, micropatterned grids without Ab or with a nonspecific Ab (gE), contained cells following a rinse (**Fig. S1**). We were also able to use this workflow to capture and position primary T-cells on EM grids micropatterned with anti-CD3 islands (**Fig. S3**).

**Fig. 1.**
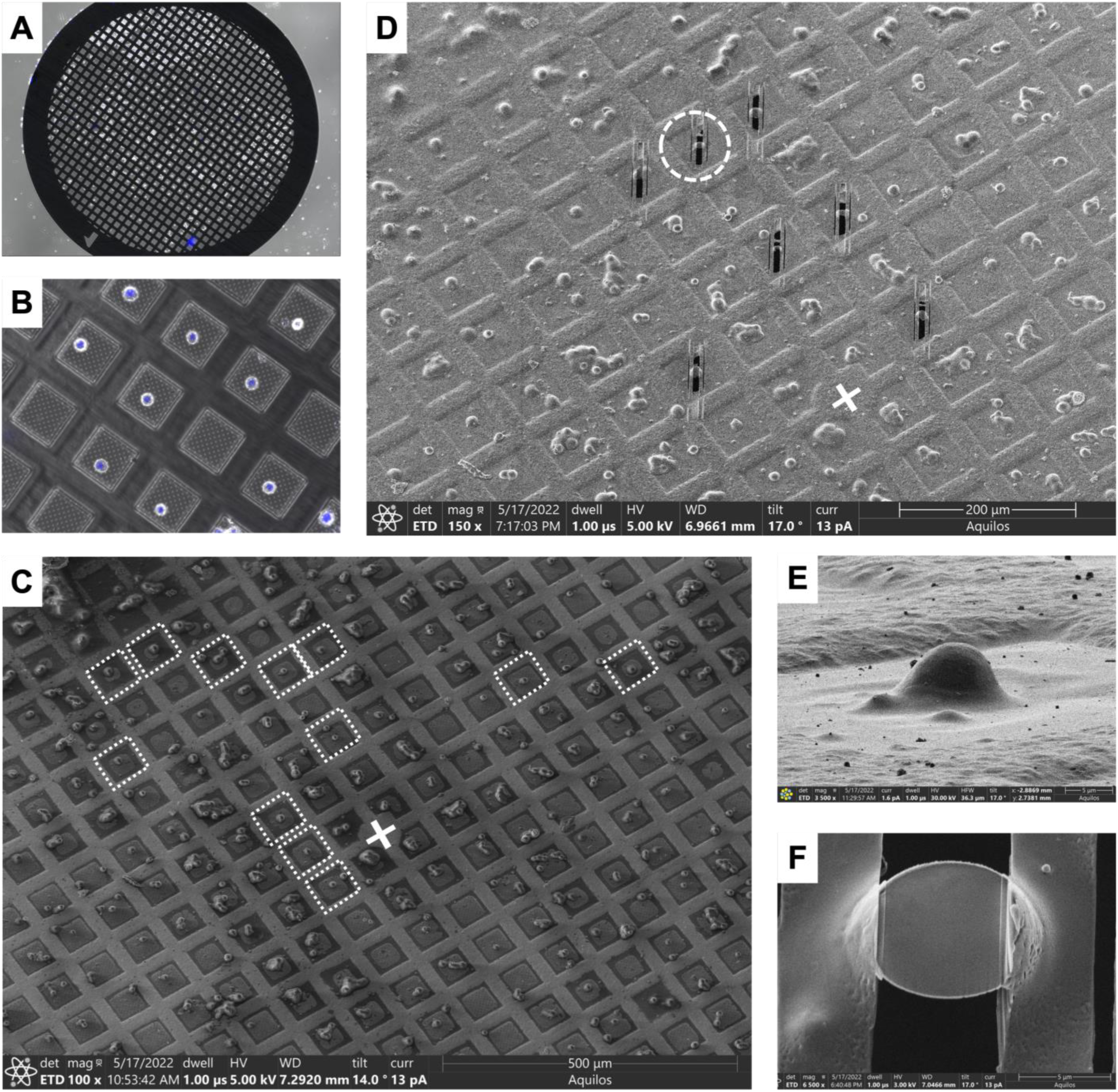
Affinity capture EM grids facilitate cryo-FIB of minimally adherent cells. A. – Fluorescence image overlaid with bright field image of 300 mesh EM grid micropatterned with 10 µm circles following antibody incubation and cell capture. **B –** Close up shows single cells centrally positioned on grid squares (nuclei are in blue). **C –** Cryo-SEM image of an EM grid with vitrified cells centrally positioned in grid squares suitable for cryo-FIB (denoted by dotted white squares). The grid center is marked with a white cross. **D –** Cryo-SEM image of the EM grid from **C** after cryo-FIB milling of seven Jurkat cells, with the grid center marked with a white cross. As is typical for cryo-FIB, all milled cells are north of the grid center. **E –** Ion beam image of a platinum-coated Jurkat cell selected for cryo-FIB milling, indicated in **D** by the white dotted circle. **F –** The lamella generated from the cell in **E** by cryo-FIB milling.

The number of Jurkat cells occupying a given grid square increased with the size of the micropatterned Ab island (**Fig. 2**). Micropatterned islands that were about half the size of Jurkat cells (i.e., 5 µm) were unable to capture the cells. Circular 10 µm islands of micropatterned anti-CD3 were ideal for vitrification and cryo-FIB/SEM of Jurkat cells (**Fig. 1C to 1F**). A typical affinity grid seeded with Jurkat cells contained more well-positioned single cells than can be milled in a typical manual cryo-FIB session (>10) (**Fig. 1C and 1D**). The target thickness (175 nm) of individually cryo-FIB milled Jurkat cells was readily achieved (**Fig. 1E and 1F**).

**Fig. 2.**
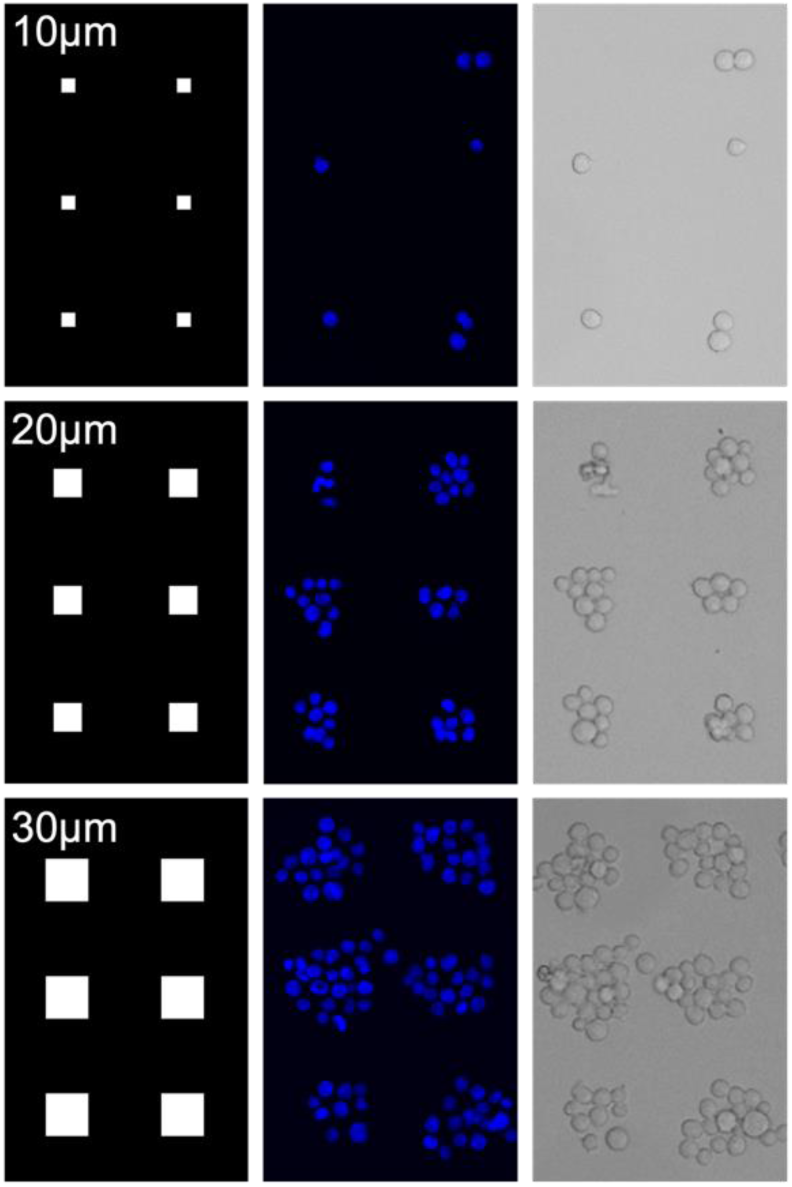
Cell occupancy increases with larger micropatterned anti-CD3 islands. Examples of micropatterned squares with side lengths ranging from 10 µm to 30 µm. Left panel: digital micropatterns designed to match the pitch of a 300 mesh EM grid. Jurkat cells attached to micropatterned glass incubated with an anti-CD3 mAb. DAPI (middle panel), bright field (right panel).

Cryo-ET of the cryo-FIB milled lamellae revealed cellular features within and external to the Jurkat cells (**Fig. 3, 4, 5, and** **S4**). Jurkat cells are designated as a T lymphoblast and their morphology, which comprises a large nucleus with very limited cytoplasm, was apparent in the montage images captured by cryo-TEM (8,700x magnification) of the lamellae (**Fig. 4**). While Jurkat cells had a smooth and round appearance when observed with light microscopy and cryo-SEM (**Fig. 1, 2, and** **3**), extracellular, filamentous structures were revealed at the periphery of the cells in the tomograms reconstructed from the cryo-ET tilt series (**Fig. 3, 4, and** **5; Movie S1 to S5**). The filaments were observed only at the trailing edge of lamellae, which captures the interface between the Jurkat cell and the SiO2 surface of the micropatterned grid (**Fig. 4, Table S1**). The filaments emanated from the plasma membrane of the cells, observed in the aligned tilt series and reconstructed tomograms (**Fig. 5; Movie S1, S2, S4 and S5**).

**Fig. 3.**
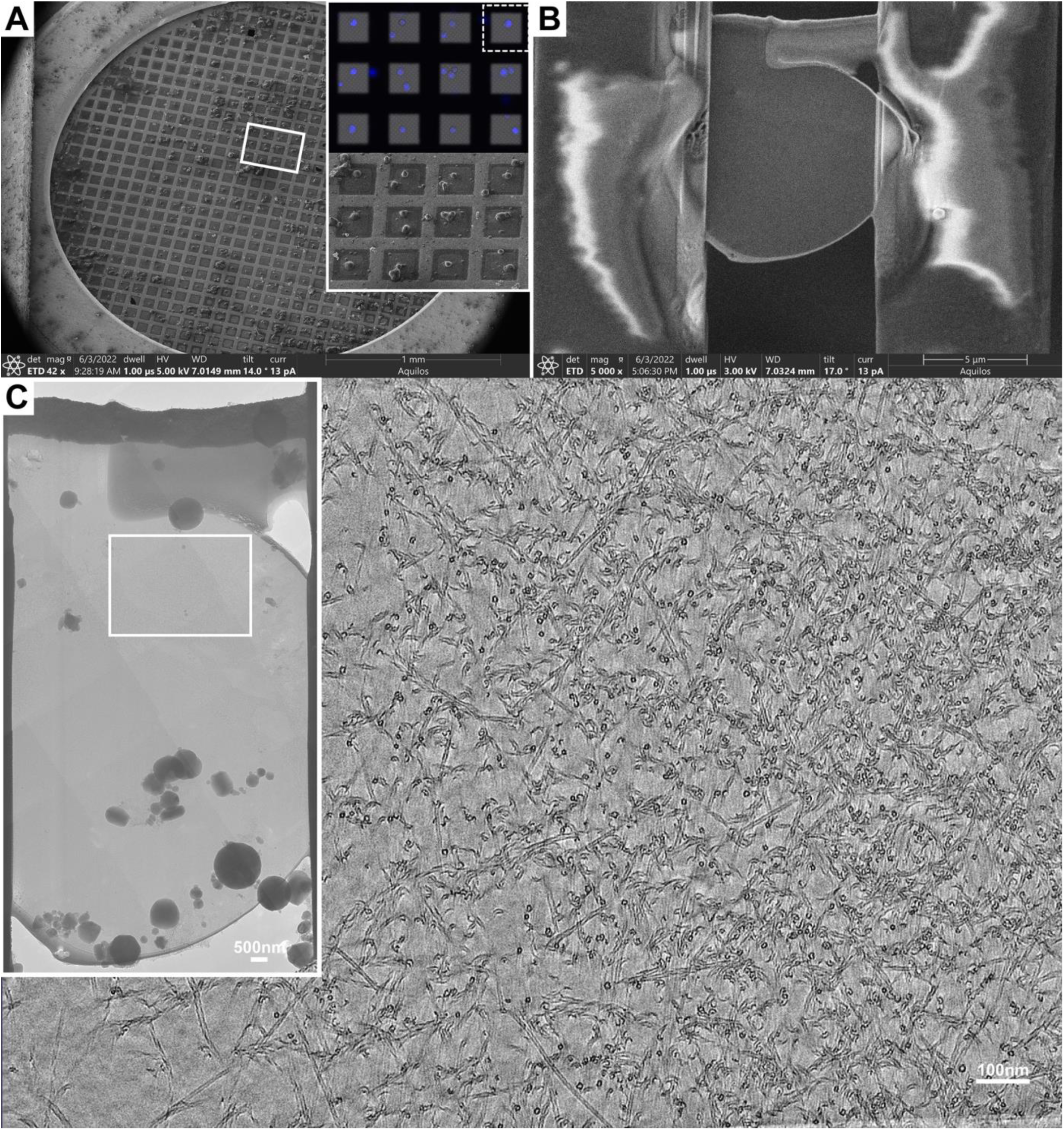
Cryo-FIB/cryo-ET identifies filaments emanating from Jurkat cells on affinity captured grids. A –. Cryo-SEM of cells vitrified on an affinity capture grid micropatterned with 10 µm circles. Inset shows live-cell fluorescence micrograph of nuclei stained with Hoechst 33342 as well as a close up on the SEM image. **B –** A lamella produced from a Jurkat cell via cryo-FIB milling with an Aquilos2. The lamella was produced from the cell highlighted by the dotted box of the inset in panel **A**. **C –** Cryo-ET of the lamella from panel **B**. The inset shows a medium magnification (8,700X) montage of the lamella. The white box highlights the area where a tilt series was collected to produce a cryo-tomogram; a single slice is shown (see Movie S4 for the entire tomogram).

**Fig. 4.**
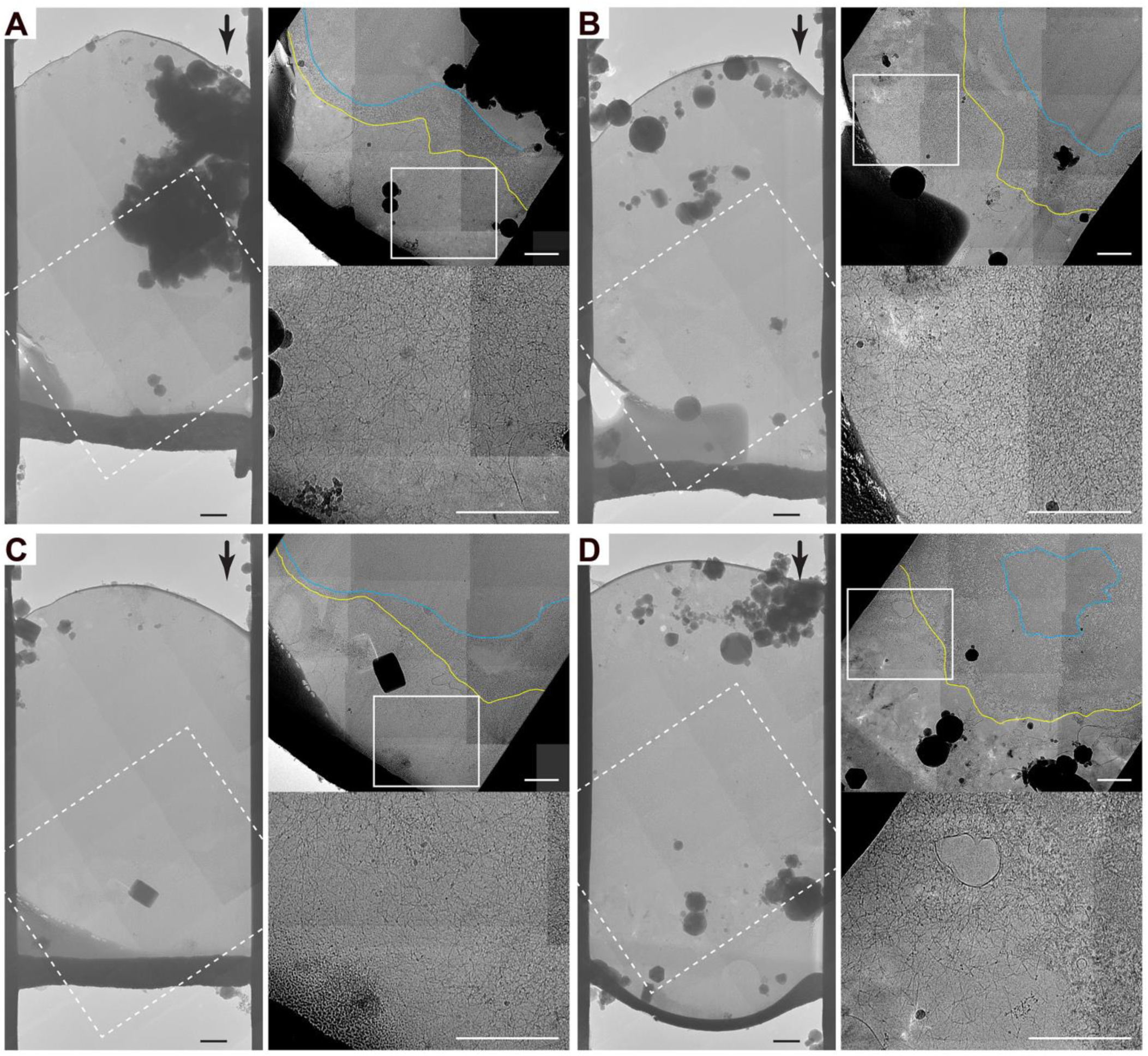
Jurkat cell extracellular filaments were localized at the SiO2 substrate-cell interface. A to D **–** Four examples of cryo-TEM montages (8,700x; 20Å/pixel) of lamellae cryo-FIB milled from micropatterned Jurkat cells. The black arrow in each left hand panel points to the thin layer of platinum coating the top of the cell and indicates the direction of cryo-FIB milling. The region where the thicker dark band below the cell meets the lighter gray region is the SiO2-Jurkat cell interface. The dotted white boxes highlight the areas shown zoomed-in in the top right image of the panel. The blue lines represent the outline of the nuclear envelopes, the yellow lines represent the outline of the plasma membrane, and the white box highlights the zoomed in area in the lower right panel revealing the intermediate filaments. The contrast for the two right hand panels in **A** to **D** was adjusted to improve visibility of the filaments. All scale bars, 1000 nm.

**Fig. 5.**
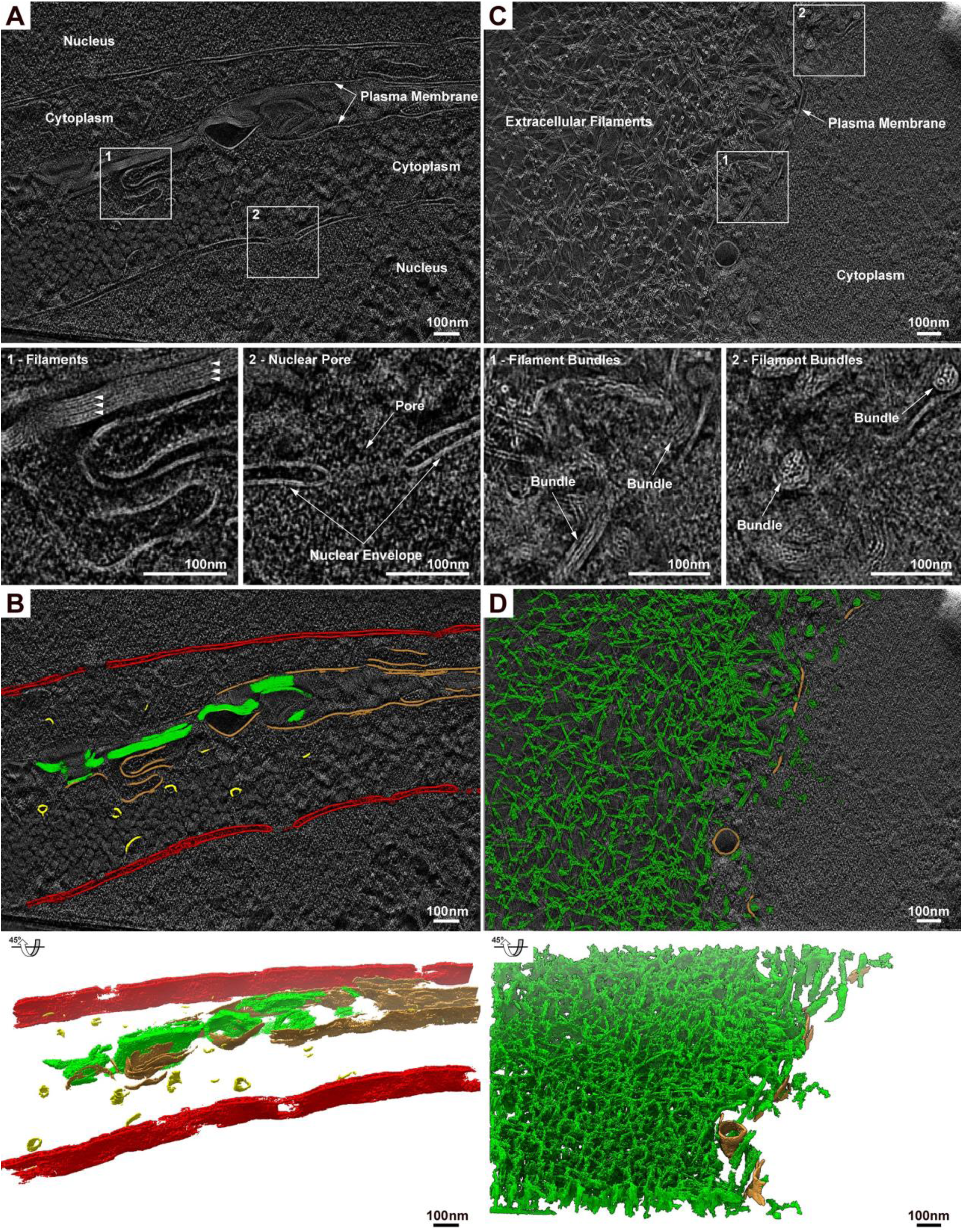
Affinity capture grids facilitate the visualization of nanoscale structures at the plasma membrane of Jurkat cells via *in situ* cryo-ET. A to D – Tomogram reconstructions and AI-based segmentation of cryo-FIB milled Jurkat cells. **A –** Cryo-tomographic slice of the region between adjacent cells showing (1) bundled filaments and (2) a nuclear pore. **C –** Cryo-tomographic slice showing extracellular filaments. In the region closer to the plasma membrane filament bundles can be seen from the side (1) and in cross section (2). **B and D –** AI-based segmentation of filaments (green), plasma membrane (brown), vesicles (yellow), and nuclear envelope (red) identified in the tomograms for panels **A** and **B**. See Movie S5 for full tomograms and Figure S4 for lower magnification images of corresponding cells and lamellae. All scale bars are 100 nm.

Within the reconstructed tomograms, the dimensions of the filaments were determined to have an external diameter of approximately 10 nm and an apparent internal diameter of approximately 3 nm, bringing them into the range of intermediate filaments (**Fig. S5**). Based on the lack of electron density, the filaments had a hollow appearance. Although the length of the filaments could not be determined from the cryo-tomograms, they are likely on the order of micrometers as they traverse the entire field of view of the tilt series and can be seen forming an extensive net-like structure in the medium magnification montages (**Fig. 3C and 4; Movie S1 to S3**). In addition, the extracellular filaments appeared to form quasi-random unconnected networks close to the cell surface but also appeared to form membranous bundles of at least 2 to 7 filaments, especially prominent in the regions between contacting cells (**Fig. 5A, 5B; Movie S1, S2 and S5; Table S2**). The shorter distance between the filaments in the bundles between cells (8.6 nm [0.124]) compared to filaments in bundles open to the extracellular environment (12.49 nm [0.622]), suggests that the structures are compressible (**Table S2**). Segmentation of the cryo-tomograms also revealed the nuclear envelope, vesicular structures, and plasma membrane (**Fig. 5B and 5D**).

The abundance of filaments within the cryo-tomograms allowed for the reconstruction of a cryo-EM map (**Fig. 6**). The architecture of the intermediate filaments, which had a diameter of approximately 100Å suggested a parallel arrangement of proteins with long, bundled alpha helices based on the recent 7.2Å cryo-EM map of polymerized vimentin intermediate filaments ^16^. The map density had a volume that could accommodate the eight alpha helix bindle of a vimentin protofibril repeating unit (**Fig. S7**). Cross- section views of the cryo-EM map revealed repetitive units of either 3 or 4 elongated structures, presumably bundled alpha helices (**Fig. 6A**). Similar cross-section densities were observed in the tomograms supporting the cross section views of the cryo-EM map (**Fig. 6A and 6C**). The map density also revealed potential interlocking sites towards the ends of the presumed helical bundles; the cryo-EM map density was diminished, indicative of flexibility between the helical bundle connections. These features are indicative of intermediate filaments, which are composed of proteins with a central alpha helical rod domain and a variable head and tail that form coiled-coiled dimers ^17^.

**Fig. 6.**
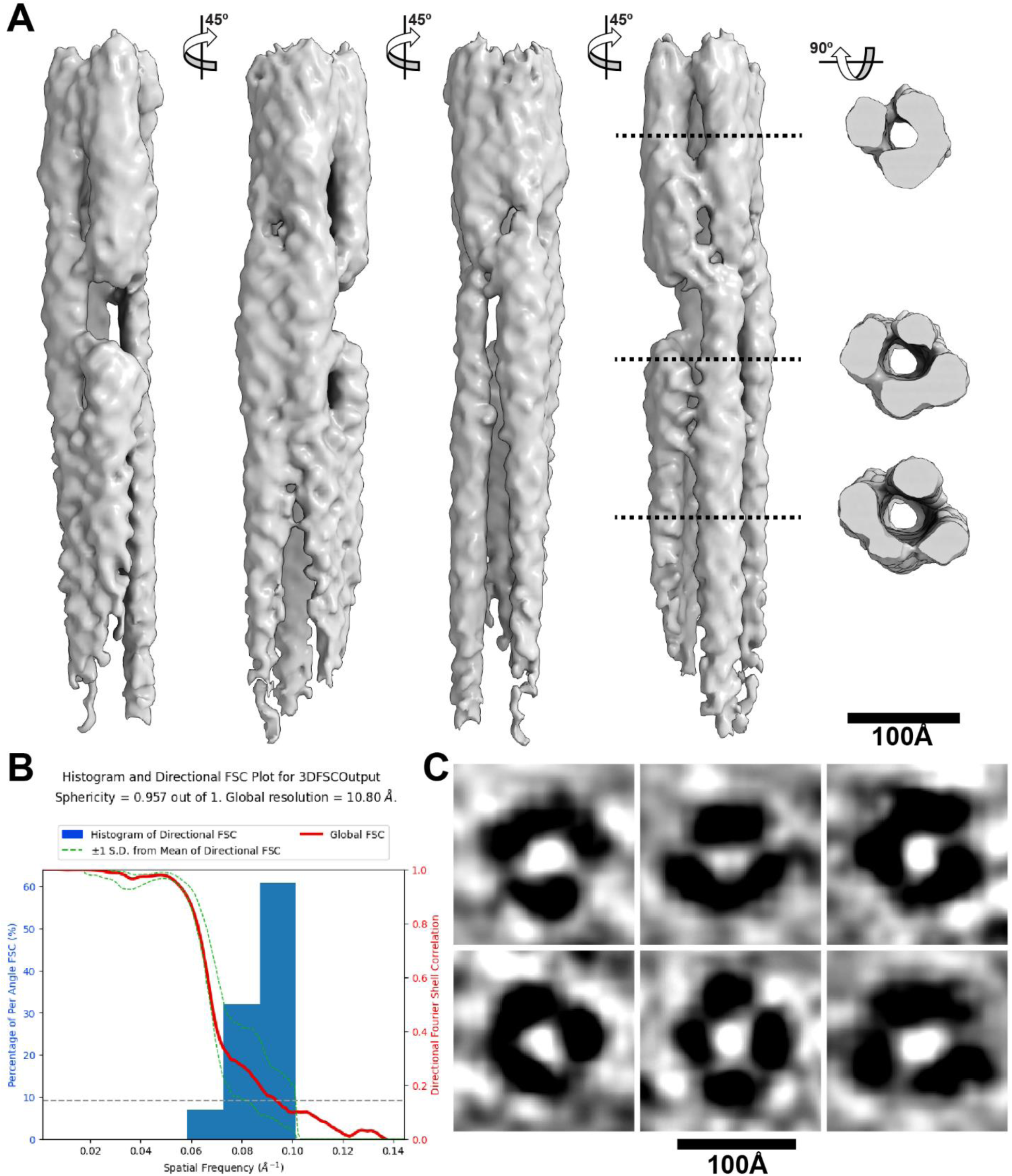
Cryo-EM map of Jurkat cell extracellular intermediate filaments. A –. Side and cross section views of the cryo-EM map of intermediate filaments reconstructed using cryoSPARC. Four views (left panels) of the cryo-EM map rotated by 45° along the Y-axis and three cross section views (far right panel) after a 90° rotation along the X-axis. **B –** A 3DFSC plot of the cryo-EM map with a reported sphericity of 0.957 out of 1 and a global resolution of 10.80Å. **C –** Representative views of individual intermediate filaments from reconstructed tomograms that reflect the cross section views of the cryo-EM map in panel A. Scale bars, 100Å.

Because the resolution of the cryo-EM map (10.8Å) was insufficient to identify the constituent proteins of the filaments, RNA-seq was performed to determine the levels of transcription from intermediate filament genes of Jurkat and control human embryonic lung fibroblast (HELF) cells. Of the 71 genes associated with intermediate filament proteins, only two type I/II (KRT1 and KRT10) and one type III (VIM) were abundantly expressed (>100 normalized reads) compared to eight type I/II/III genes for the HELF controls (**Table S3**). Vimentin (VIM) was detected on the external surface of Jurkat cells by flow cytometry of non-permeabilised (surface) and permeabilised (total) cells (**Fig. S8 and S9**). Compared to no antibody or isotype controls, vimentin was detected in almost all (>98%) permeabilised and non- permeabilised Jurkat cells. The levels of total and cell surface vimentin were not altered by incubating Jurkat cells with an isotype control, anti-CD4 or anti-CD3 antibodies prior to fixation. These data suggest that the extracellular filaments observed by cryo-ET are vimentin intermediate filaments.

## DISCUSSION

We have demonstrated an affinity capture system to accurately and repeatedly position lymphocytes on EM grids for cryo-FIB milling and cryo-ET. T-cells have been observed in the literature to form stable attachments to micropatterned islands of CD3 antibodies on glass surfaces ^18^, which is likely the mechanism underlying affinity capture on EM grids. Affinity capture grids facilitated our observation of nanoscale filaments on the surface of Jurkat cells in several of our tomograms (**Figs. 3, 4 and 5; Movies S1 to S3**). The dimension of the filaments measured from our tomography data (10 nm, **Fig. S5**) was uniform and consistent with intermediate filaments, such as keratin, which have recently been observed in the extracellular space ^19,20^. Because the filaments were outside of the cell plasma membrane, we considered the possibility of it being a glycocalyx, which is known to have a filamentous architecture, but in contrast to the filamentous unconnected network we observed, the glycocalyx is heavily interconnected with a wide distribution of filament diameters (3 to 15 nm)^21,22^.

The elongated structures seen in the cryo-EM map (**Fig. 6** **and S7**) together with the high levels of expression seen in the RNA sequencing (**Table S3**), and detection of vimentin on extracellular surfaces by flow cytometry (**Fig. S8 and S9**), support vimentin as the major constituent of the filamentous networks observed in the Jurkat cell extracellular space. Canonically, intracellular vimentin intermediate filaments are known to provide structural support to lymphocytes ^23,24^, but vimentin also plays crucial roles in lymphocyte migration and attachment to the vascular endothelium ^25–28^. Vimentin has also been detected on the extracellular side of the plasma membrane in viable malignant lymphocytes, normal activated T- cells, and apoptotic T-cells ^29^, although the role of extracellular vimentin, how it is secreted, and what domains of vimentin are expressed on the surfaces of different cell types (e.g., lymphocytes vs. endothelial cells) remain unclear ^28,30,31^. The captured Jurkat cells observed in the present study might have been activated by the anti-CD3 antibody and are a malignant cell line such that the presence of extracellular vimentin would be consistent with previous studies ^29^. Extracellular vimentin in human atherosclerotic tissue lesions was detected in areas of inflammation and in the necrotic core ^32^. In addition, it is possible that fixation, as performed in this study prior to plunge-freezing, provided a stressful condition that influenced the presence of extracellular vimentin. Of note, prior studies reporting on extracellular vimentin used chemical fixation ^29,33,34^. The functional implications of extracellular vimentin across cell types have recently been reviewed by Suprewicz, and Thalla and Lautenschlager ^35,36^.

Vimentin monomers assemble into parallel homodimers that form higher order structures of anti- parallel adjacent dimers then tetramers to form the underlying asymmetric unit of a vimentin intermediate filament protofibril ^16^. Three tetramers are necessary for the complete assembly of a vimentin intermediate filament protofibril with eight polypeptide chains at the core of the repetitive unit. The cryo-EM map of the Jurkat cell extracellular intermediate filaments is highly suggestive of such a higher order vimentin complex given the long densities that could accommodate the eight alpha helices of the vimentin protofibril repetitive unit (**Fig. S7**). Intriguingly, the extracellular filaments in the present study had an electron transparent lumen and appeared to be composed of 4 protofibrils, which was in contrast to the exogenously expressed, polymerized human vimentin ^16^. This recent study also reported that intracellular vimentin intermediate filaments of mouse embryonic fibroblasts assembled into higher order structures and supports the helical assembly of 5 spring-like protofibrils ^16^ . However, the 4 protofibril architecture of the extracellular vimentin from Jurkat cells suggests that the role of vimentin intermediate filaments is context specific and cell type dependent. Furthermore, it remains unclear how these long, apparently membrane-less projections, emerge from the cell.

To the best of our knowledge, this is the first time networks of native intermediate filaments have been observed at the nanoscale in the extracellular space. Jurkat cells have been observed to shed microvillus membrane particles on anti-CD3 treated surfaces using total internal reflection fluorescence microscopy (TIRF)^14^, but the filaments that appear in our cryo-tomograms (**Fig. 3C and 4D**) are too small (10 nm) in diameter) and very densely packed to be observed by fluorescence microscopy. The filaments were also not observed by conventional transmission and scanning EM (TEM and SEM) imaging of the Jurkat cell surface^14^. While these EM modalities can provide higher resolution than light microscopy, in contrast to *in situ* cryo-ET, they require preparation steps that would likely cause the collapse of delicate structures such as networks of filaments on the cell surface. Specifically, for conventional SEM of cells, preparation steps include fixation, dehydration, and the deposition of a conductive metal layer. For conventional TEM, preparation of cells includes fixation, dehydration, resin embedding, staining, and sectioning. Cryo-ET overcomes the limitations of these destructive processing methods, providing the highest resolution imaging of cells preserved in a hydrated, near-native state. Cryo-FIB allows for the generation of nearly distortion-free thin sections of vitrified cells ^37–39^. In our cryo-FIB/cryo-ET study, preservation of nanoscale biological structures through vitrification converged with three-dimensional image processing techniques to provide an unprecedented view of the nanoscale environment of the Jurkat cell surface. Other recent studies have demonstrated the power of the cryo-FIB/cryo-ET platform to reveal additional complexity to biological structures such as microvilli despite decades of structural analysis by conventional EM ^40^.

The affinity capture system improves sample usability, reducing the cost and preparation time required to prepare cellular samples for cryo-FIB/cryo-ET. Beyond positioning cells and limiting the number of cells that attach to a given region of the substrate, the use of micropatterned islands functionalized with antibodies, ligands, and ECM proteins can potentially be extended to select for and capture cell types of interest from diverse cell populations. Alternatively, two different antibodies can potentially be patterned to capture different cell types within the same grid square for co-culture studies. We envision the workflow presented here facilitating future analysis of the nanoscale structures involved in T-cell activation. Patterns comprised of different ligands can potentially be used to study how the spatial organization of signaling complexes impacts the structures involved in cell communication ^18^, such as receptor arrangement ^41^. An extension of this idea is to facilitate studies of the nanoscale architectures involved in signaling events or the growth and structure of nanoscale bridges (i.e., tunneling nanotubes) by using micropatterning to control the distance between captured cells ^42,43^.

Appropriately packaged affinity capture EM grids could assist non-experts (e.g., clinicians lacking EM experience) with preparing and shipping clinically-relevant cells to cryo-EM facilities for cryo-ET imaging. Packaging methods currently include the use of 3D printed grid holders ^44^ and silicone wells in glass-bottom dishes that exploit the surface tension of liquid droplets contained to the wells to immobilize EM grids ^45^. Devising storage methods that increase the shelf life of micropatterned EM grids is an important future area of research toward increasing access to affinity capture EM grids.

## METHODS

### EM Grid Micropatterning

To accommodate single Jurkat cells, squares and circles varying from 5 to 30 µm across were micropatterned on 300 mesh gold grids coated with a thin perforated layer of silicon dioxide (Quantifoil R1/4). The grids were exposed to atmospheric plasma (PE-50, Plasma Etch Inc., Carson City, NV, USA) at 30 W for 12 seconds to render the surface hydrophilic, immediately immersed in 0.01% poly-l-lysine (PLL, Sigma cat # p4707) and incubated in PLL overnight at 4 °C, and then rinsed and incubated in a solution of 100 mg/mL mPEGSuccinimidyl Valerate, MW 5,000 (PEG-SVA, Laysan Bio Inc) in 0.1M HEPES (pH 8.5) the following day for 1 hour at room temperature. Following PEG coating, 3 *μ*L of a 1:6 solution of PLPP photoinitiator (Alveole) in ethanol was added to the surface of each grid and allowed to air dry while being protected from light. The grids were then UV exposed according to digitally generated micropatterns at 50 mJ/mm^2^ using the Primo photo-micropatterning system (Alveole) to selectively degrade the PEG layer on the EM grids and rinsed in deionized water. A detailed protocol for micropatterning EM grids using this method has been made publicly available by the authors on the protocol sharing platform protocols.io ^45^.

Following micropatterning of the grids and sterilization in 70% ethanol (see aforementioned protocol), the grids were incubated in human anti-CD3 (clone UCHT1; Biolegend Cat# 300402) antibody overnight. Control conditions included a set of micropatterned grids that were not incubated in antibody (No Ab) and another set of micropatterned grids that were treated with monoclonal antibodies to VZV glycoprotein E (gE; EMD Millipore Cat# MAB8612).

For T-cells, 200 mesh gold Quantifoil holey carbon R2/2 grids were micropatterned followed the same protocol, but with micropatterned circles that were 7.5 and 15 µm in diameter.

### Jurkat Cell Culture and Seeding

Jurkat cells (Jurkat, Clone E6-1, ATCC) were maintained following manufacturer instructions. A volume of 10 µL of a 7x10^6^ cells/mL suspension was seeded on the micropatterned grids and allowed to settle for two hours. The grids were then transferred to fresh 35 mm dishes with 10 mm 1.5 cover glass (MatTek) and washed extensively with media (RPMI 1640 + 10% FBS + 1% pen/strep). The grids were then washed once with 2 mL phosphate-buffered saline (PBS, Gibco), fixed in 1 mL 4% paraformaldehyde (PFA), and washed twice with 2 mL PBS. Prior to imaging, the PBS was aspirated and replaced with 2 mL PBS plus 1:1000 Hoechst 33342 to stain the cell nuclei. Images were captured with a BZ-X710 microscope (Keyence). Brightness and contrast were adjusted to improve visibility (**Fig. 2**).

### T-cell Culture and Seeding

Human T-cells (gift from Robbie Majzner’s lab at Stanford University) were purified from isolated peripheral blood mononuclear cells as in Tousley et al. ^46^. They were maintained in AIM-V complete media (Gibco) supplemented with 5% fetal bovine serum (FBS) and IL-2 (2.18 IU/ng, Peprotech). To seed the cells on the micropatterned grids, 25 µL of a 4x10^5^ cell/mL suspension was added to each grid several times until approximately 2 to 3 cells were observed above or on each grid square. Images of live T-cells on micropatterned Au 200 mesh holey carbon coated EM grids were captured after 5.5 hours on an inverted Nikon Ti-E microscope (Nikon, Minato, Tokyo, Japan) equipped with a Heliophor light engine (89 North) and an Andor sCMOS Neo camera using a 20x Plan Apo Lambda air objective lens and a 60x Plan Apo Lambda oil objective lens. The cells were vitrified by plunge freezing.

### Vitrification and Cryo-FIB

Samples were vitrified in a Leica EM GP2 plunge freezer after blotting from the back side of the EM grids for 9 seconds. The vitrified grids were clipped into autogrids before being loaded into an Aquilos 2 (Thermo Fisher) for cryo-FIB milling. Lamella were cryo-FIB milled at a 10 degree milling angle first to a 5 µm thickness with a current of 1 nA. Microexpansion joints were cryo-FIB milled to prevent bending and breaking of the cryo-lamellae ^47^. Sequential milling was performed at 300 pA to 3 µm thickness, at 100 pA to 1µm thickness, and at 50 pA to 500 nm thickness. A final polish was performed with a 175 nm distance between rectangular patterns at a current of 30 pA.

### Cryo-ET Data Acquisition and Tomogram Reconstruction

Samples were loaded into an FEI Krios G2 transmission electron microscope (TEM) equipped with a bioquantum energy filter and a K3 direct electron detector (Gatan). SerialEM software ^48^ was used to operate the TEM at 300 kV in low-dose mode and acquire tilt series at a magnification 26,000x (3.465 Å/pixel). Tilt series were acquired with 2*°*steps between -60*°* and +60*°* and a defocus of -4 µm. Additional data collection parameters are in Supplemental Table 4. Motion correction and tilt series stack generation was performed with Warp software ^49^ and 3D reconstructions were calculated using the weighted-back projection using IMOD ^50^. AI segmentation of tomograms was performed using the Dragonfly (Version 2022.2.0.1367; Object Research Systems) Deep Learning Tool. UCSF Chimera was used for visualization of structures ^51^.

Movies were generated from individual frames using ffmpeg^52^ available from http://ffmpeg.org/.

### Cryo-ET Map Reconstruction of Extracellular Intermediate Filaments

CryoSPARC^53^ was used to generate a reconstruction of the intermediate mediate filaments using the helical processing pipeline (**Fig. S6**). CryoSPARC does not currently have a dedicated cryo-ET processing pipeline. However, due to the limited number of tilt series (three) for intermediate filaments at the periphery of the Jurkat cells, individual movie stacks from each tilt of the cryo-ET data acquisition were imported (Import Movies) into a CryoSPARC project with the accumulated dose information for each movie stack. Movies where the edges of the lamella wall entered the field of view were removed, leaving a total of 110 movies. CryoSPARC’s patch motion correction and CTF estimation was performed on the 110 movies (micrographs). To generate a 2D class average for template picking, 238 particles were manually picked from 12 micrographs. These particles were subjected to 2D classification into 4 classes with the ‘align filament classes vertically’ option checked; three classes contained the majority of particles (237) and were used for filament tracing with the following parameters; Filament diameter (100Å), Separation distance between segments (25Å), Minimum filament length to consider (400Å), Angular sampling (5°). The 250,941 particles generated were extracted from the 110 micrographs with a box size of 120 pixels, which were used for a further round of 2D classification into 50 classes. Of these, 3 classes (46,827) were used for an additional round of filament tracing, extraction of particles from the micrographs with a box size of 240 pixels to increase the length of the filaments captured, and 2D classification of the newly extracted particles (382,315) into 50 classes. Of these, 85,475 particles from 5 classes were used for *de novo* reconstruction using ‘Helix refine’ then subjected to 3D classification. One class containing 83,607 particles was further processed to remove duplicate particles. Helix refine was performed on the remaining 35,789 particles yielding a 13.0Å cryo-ET map as determined by FSC estimation with loose masking. To quantitatively assess directional resolution anisotropy of the cryo-ET map, 3DFSC^54^ was used with the half maps and the refinement mask from ‘Helix refine’ used as input.

### RNA-seq of Jurkat and HELF Cells

Jurkat and HELF cells were lysed with buffer RLT Plus following the manufacturer’s instructions (Qiagen). RNA was purified from lysates using an RNeasy Plus Mini Kit (Qiagen). RNAseq libraries and sequencing (RNA-seq) was performed by Medgenome. FASTQ files generated from the libraries were assessed and aligned to the human genome (hg19) using the STAR aligner in RNAdetector ^55^. To assess expression levels for from intermediate filament genes, normalized reads produced by RNAdector were used.

### Flow Cytometry of Jurkat Cells

Jurkat cells were incubated with either CD4-FITC (clone: RPA-T4; Biolegend), CD3-FITC (clone: OKT3; Biolegend), or isotype-FITC (clone: MOPC-173; Biolegend) antibodies (1 µl per 2 x10^5^ cells) or mock treated at 37 °C for 1 hour then fixed with 4% paraformaldehyde. All samples were blocked with Human TruStain FcX (Fc Receptor Blocking Solution; Biolegend) on ice for 10 minutes before immunostaining with vimentin rabbit mAb conjugated with Alexa fluor 647 (clone: D21H3; Cell Signaling) or isotype control rabbit mAb conjugated with Alexa fluor 647 (clone: DA1E; Cell Signaling) on ice for 30 minutes, washed twice with ice-cold PBS supplemented with 0.5% BSA (Bovine serum albumin; Jackson ImmunoResearch) and 0.02% sodium azide (Sigma), then resuspended in ice-cold PBS supplemented with 2.5% FBS, 2 mM EDTA (Fisher scientific), and 0.2 µg/ml DAPI (Thermo Fisher Scientific). Stained cells were analyzed using Agilent Quanteon flow cytometer (Agilent), color compensation was performed using compensation beads (Biolegend) only, compensation beads with CD3-FITC antibody, compensation beads with vimentin Alexa fluor 647 antibody, and Live:Dead (1:1) with DAPI stain. Data were processed with FlowJo (TreeStar) to determine the percentage of live cells (DAPI signal), percentage of cells expressing vimentin (Alexa fluor 647 signal) and percentage of cells expressing CD4 or CD3 (FITC signal).

## DATA AVAILABILITY

Data generated and/or analyzed during the current study are available in the paper or are appended as supplementary data, and data that support this study are available from the authors upon request. The cryo-ET map has been deposited in the Electron Microscopy Data Bank (EMDB) with accession code EMD-43978 and the original movie files, tilt series, and tomograms have been deposited in the Electron Microscopy Public Image Archive (EMPIAR) with accession code EMPIAR-12110. All primary data will be provided by the corresponding author upon request.

## ACKNOWLEDGEMENTS

This work was supported by NIH R01 AI20459 (SO), R21 AI159375 (SO), R35 GM130332 (ARD), and a Diane and Guilford Glazer Foundation Faculty Fellowship (LE). We thank Dr. Eva de la Serna (Dunn lab) for culturing T-cells on EM grids and Maria Caterina Rotiroti and the Majzner lab at Stanford for providing the CAR T-cells used in this study. We thank Dr. Wah Chiu (Stanford) and Dr. Matthew J. Paszek (Cornell) for helpful discussion and Dr. Lydia-Marie Joubert for providing cryo-FIB training, experimental support, and helpful feedback. We thank the Stanford CryoElectron Microscopy Center (cEMc) for providing access to a Krios cryo-TEM for data collection. Cryo-FIB for this work was performed at the Stanford-SLAC CryoET Specimen Preparation Center (SCSC), which is supported by the National Institutes of Health Common Fund’s Transformative High Resolution Cryoelectron Microscopy program (U24GM139166). The content is solely the responsibility of the authors and does not necessarily represent the official views of the National Institutes of Health. The authors declare no competing financial interests.

## AUTHOR CONTRIBUTIONS

LE: Conceptualization, Methodology, Validation, Investigation, Writing - Original Draft, Visualization, Supervision, Project administration. MZ: Investigation, Writing - Review & Editing. MZ: Investigation. ARD: Resources, Writing - Review & Editing, Supervision, Funding acquisition. SO: Conceptualization, Methodology, Software, Validation, Formal analysis, Investigation, Resources, Data Curation, Writing - Original Draft, Visualization, Supervision, Project administration, Funding acquisition.

