## Supplemental Figures for "Extracellular filaments revealed by affinity capture cryo-electron tomography"

### Supplemental Information

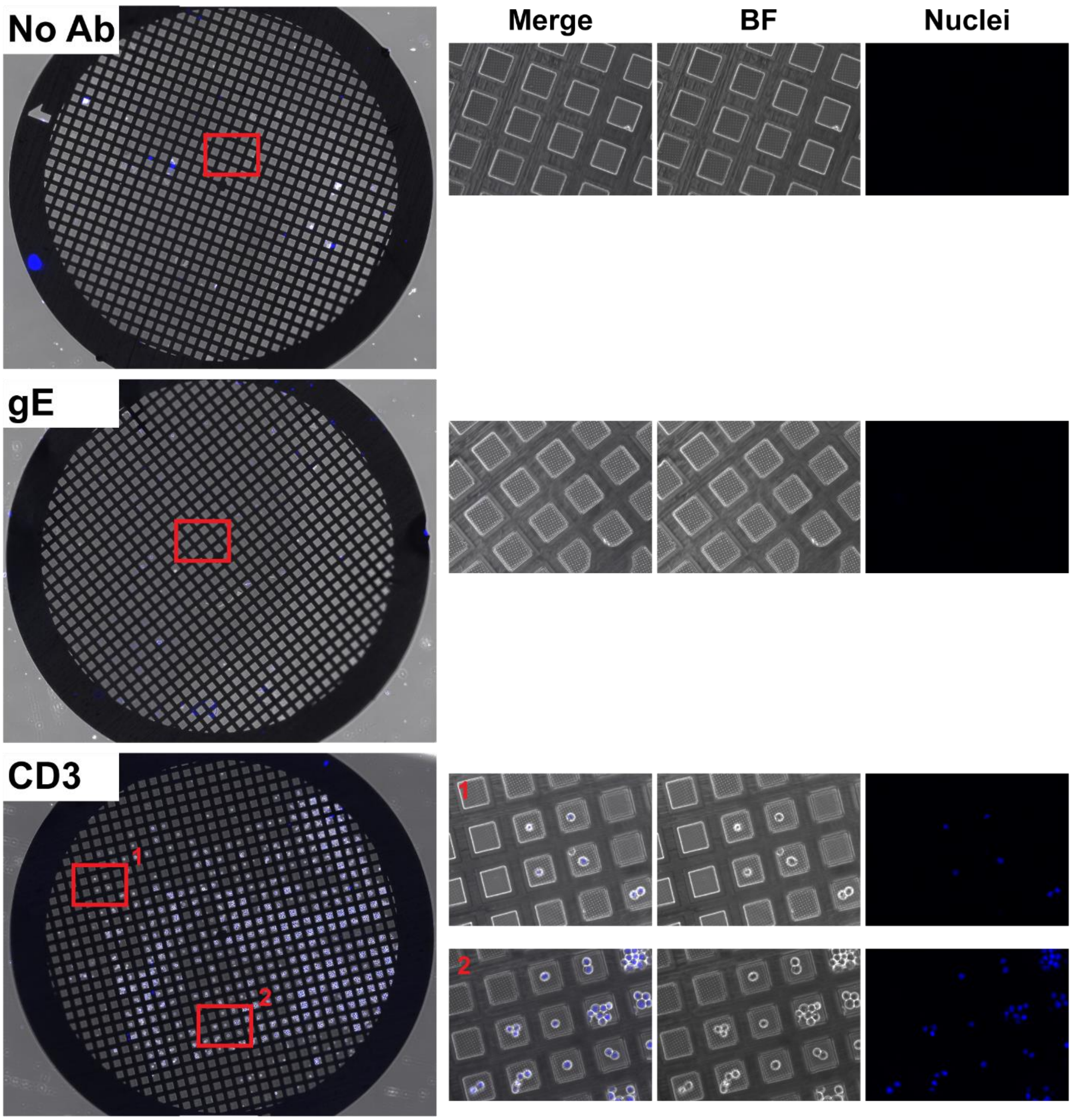


**Figure S1. Affinity capture of Jurkat cells on micropatterned electron microscopy grids.** Au 300 mesh $SiO_2$ electron microscopy grids micropatterned with 10 µm circles in the center of grid squares were either untreated (No Ab) or treated with monoclonal antibodies to VZV glycoprotein E (gE) or human CD3 (CD3). Jurkat cells (10 µl at $7x10^6$ cells/ml) were allowed to settle for two hours on the grids. The grids were transferred to fresh 35 mm dishes with 10 mm 1.5 cover glass (MatTek) then washed extensively with media (RPMI 1640 + 10 FBS + pen/strep). The grids were washed once with 2 ml PBS, fixed with 1 ml 4% PFA, then washed twice with 2 ml PBS. PBS was aspirated and replaced with 2 ml PBS plus 1:1000 Hoechst 33342 to stain cell nuclei. Images were captured with a BZ-X710 microscope (Keyence). Montaged images (left panels) of the grids were captured with a 10x objective (merge of bright field and nuclei). Zoomed-in images (right panels; locations on the grids are indicated by the red rectangles on the montaged grid images) were captured with a 40x objective.

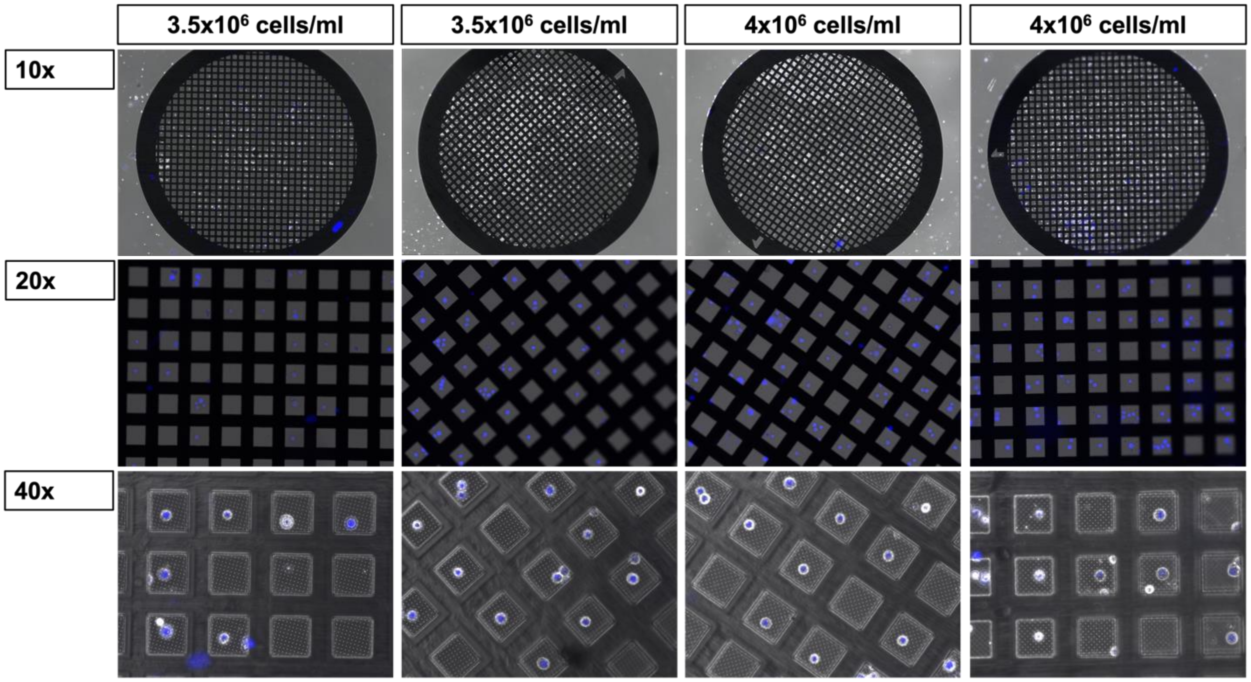


**Figure S2. Affinity capture of Jurkat cells on micropatterned electron microscopy grids.** Au 300 mesh $SiO_2$ electron microscopy grids micropatterned with 10 µm circles in the centers of grid squares with cells seeded at two different concentrations.

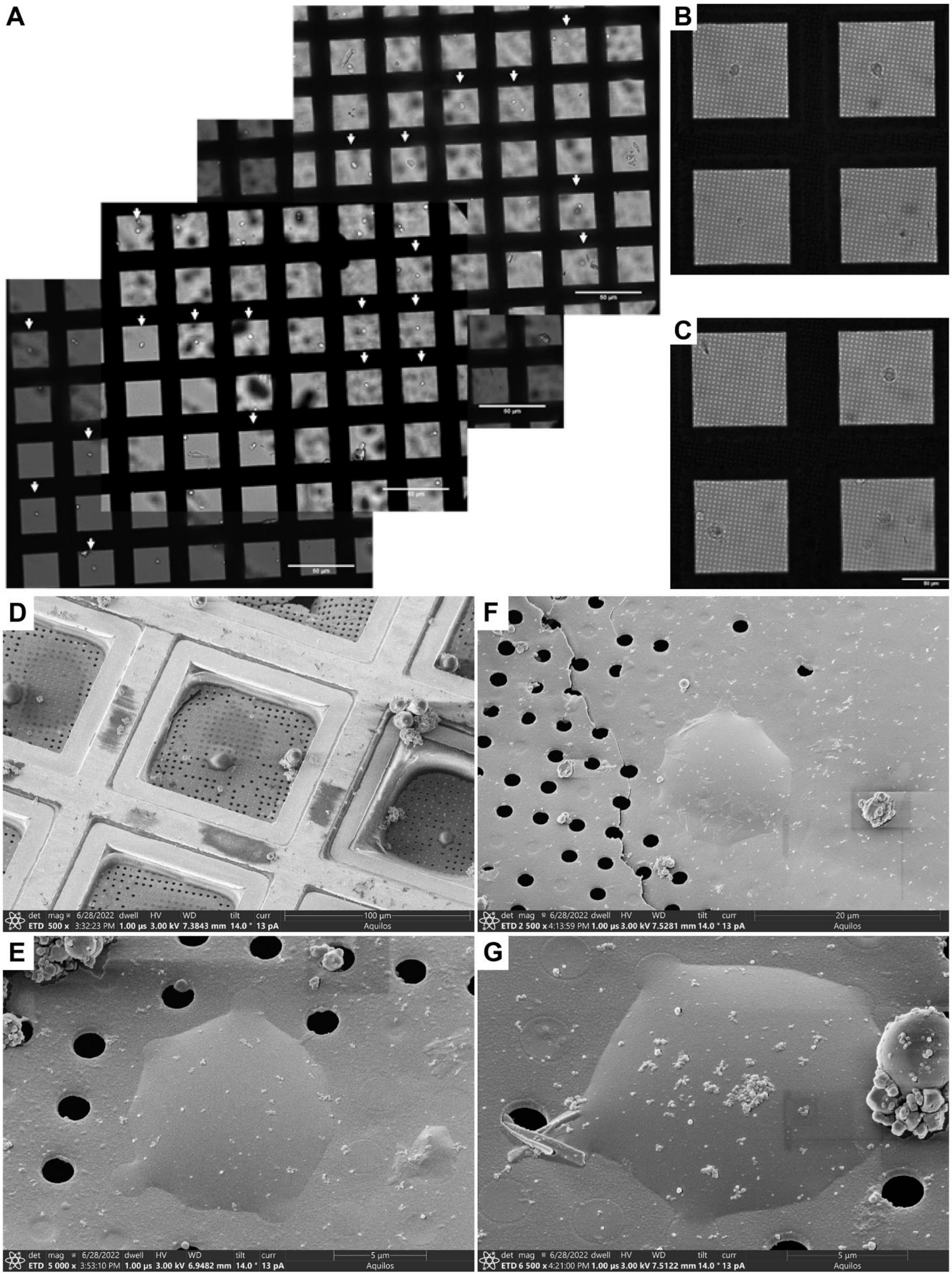
A
B
C
D
E
F
G
50 µm
50 µm
50 µm
50 µm
50 µm
det mag 6/28/2022 dwell HV WD tilt curr 100 µm
ETD 500 x 3:32:23 PM 1.00 µs 3.00 kV 7.3843 mm 14.0 ° 13 pA Aquilos
det mag 6/28/2022 dwell HV WD tilt curr 20 µm
ETD 2 500 x 4:13:59 PM 1.00 µs 3.00 kV 7.5281 mm 14.0 ° 13 pA Aquilos
det mag 6/28/2022 dwell HV WD tilt curr 5 µm
ETD 5 000 x 3:53:10 PM 1.00 µs 3.00 kV 6.9482 mm 14.0 ° 13 pA Aquilos
det mag 6/28/2022 dwell HV WD tilt curr 5 µm
ETD 6 500 x 4:21:00 PM 1.00 µs 3.00 kV 7.5122 mm 14.0 ° 13 pA Aquilos

**Figure S3. Affinity capture of T-cells on Au 200 mesh holey carbon (Quantifoil R 2/2) EM grids micropatterned with 15 µm circles of anti-CD3.** No cells were found attached to EM grids micropatterned with 7.5 µm circles. **A** – A montage image was created from bright field images taken of live, unfixed cells at a magnification of 20x. At least 19 well-positioned single cells can be counted on a single micropatterned EM grid (indicated by white arrows). **B to C** – Live cell images acquired with a 60x oil objective. **D to G** – Cryo-SEM of vitrified T-cells captured in central positions on grid squares.

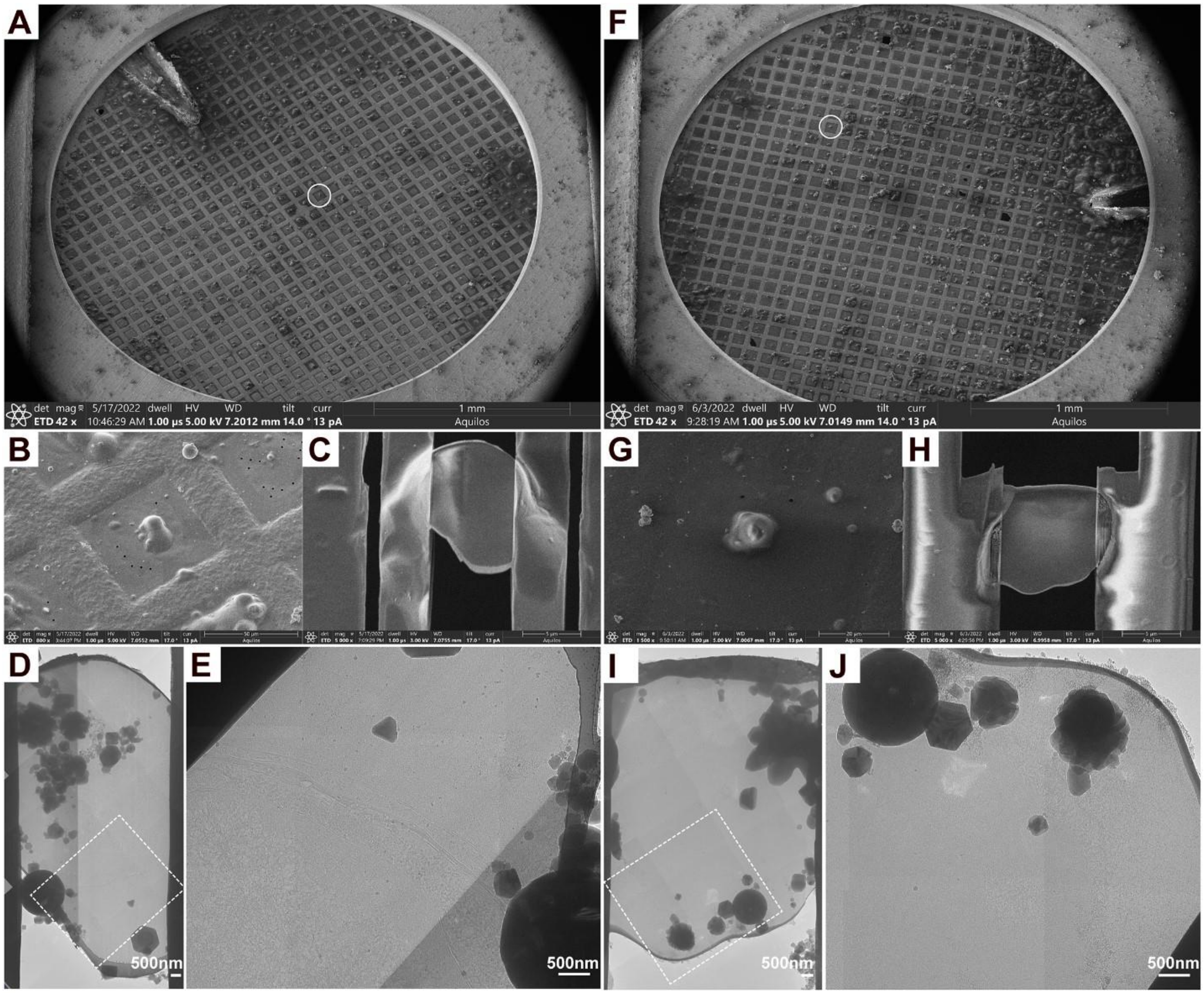


**Figure S4. Cryo-FIB/SEM milling and cryo-EM of Jurkat cells used for cryo-ET and tomogram reconstruction. A to J** – Cryo-SEM of grids (**A and F**), individual grid squares (**B and G**), and cryo-FIB milled Jurkat cells (**C and H**). Cryo-EM was used to capture medium magnification montages (**D, E, I and J**) to identify regions within the lamellae (dotted boxes) for cryo-ET tilt series (±60°) collection. The lamella depicted in A-E and F-J are those used to generate tomogram reconstructions and segmentations in Fig. 4A and B, and C and D, respectively.

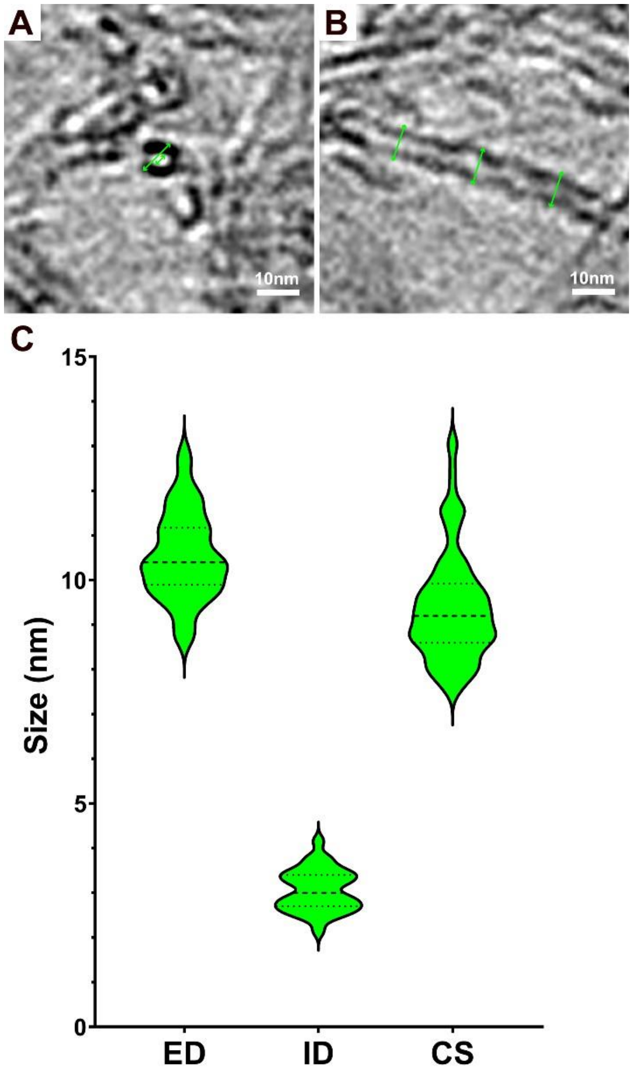


**Figure S5. Dimensions of extracellular filaments produced by Jurkat cells. A and B -** Cross sections of the intermediate filaments; transverse (**A**) and median (**B**). The green arrows show the dimensions measured; A – External (ED) and internal (ID) diameters, B – Cross section (CS). **C** – Violin plots for the dimensions of the measured filaments. For ED and ID n = 60 for each dimension across three tomograms. For CS n = 90; 30 filaments across three tomograms measured in three places. The dashed lines show the median values and the dotted lines show the 25$^{th}$ and 75$^{th}$ percentiles.

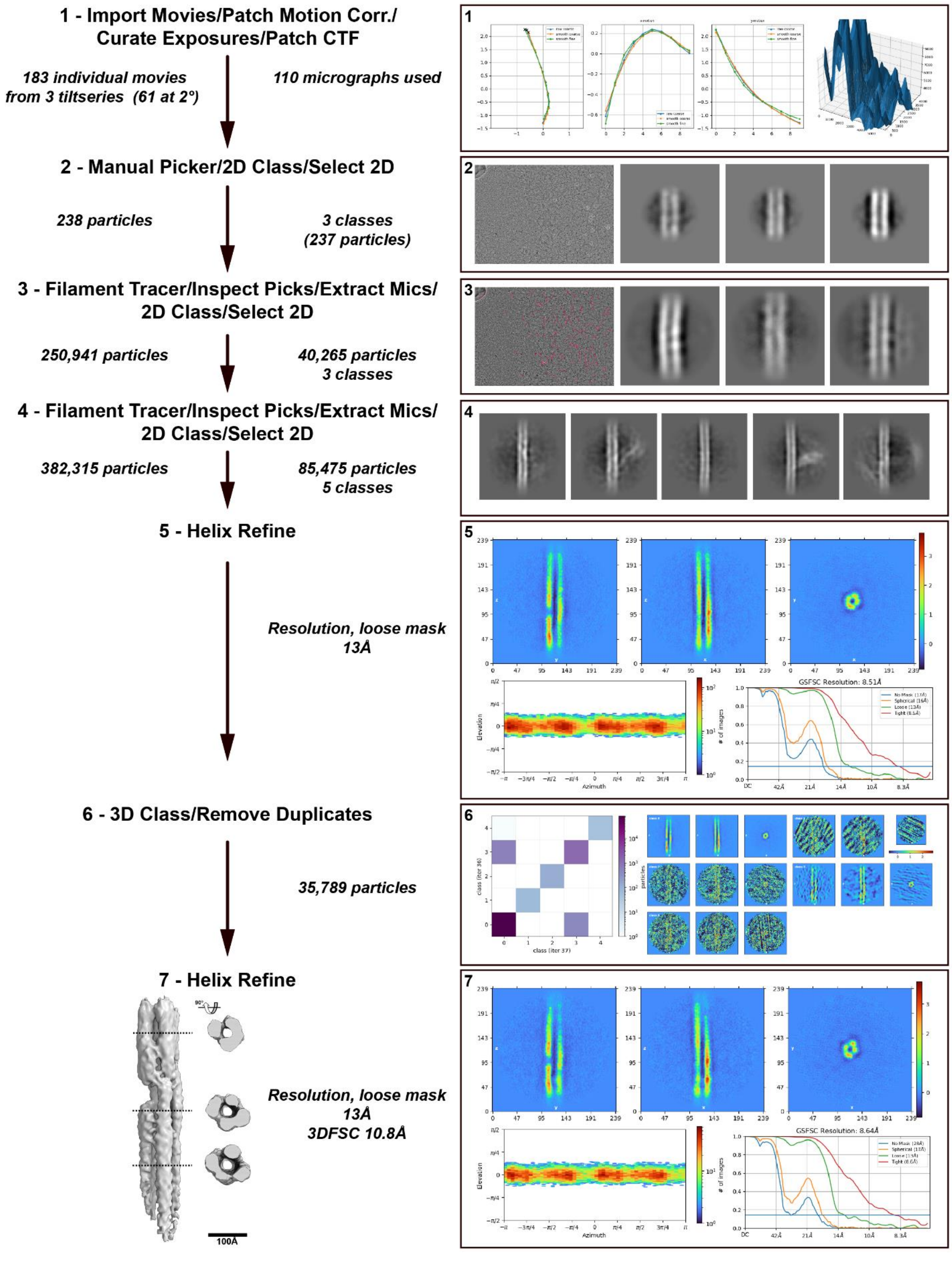

1 - Import Movies/Patch Motion Corr./
Curate Exposures/Patch CTF
183 individual movies
from 3 tiltseries (61 at 2°)
110 micrographs used
1
2 - Manual Picker/2D Class/Select 2D
238 particles
3 classes
(237 particles)
2
3 - Filament Tracer/Inspect Picks/Extract Mics/
2D Class/Select 2D
250,941 particles
40,265 particles
3 classes
3
4 - Filament Tracer/Inspect Picks/Extract Mics/
2D Class/Select 2D
382,315 particles
85,475 particles
5 classes
4
5 - Helix Refine
Resolution, loose mask
13Å
5
GSFSC Resolution: 8.51Å
Elevation
Azimuth
6 - 3D Class/Remove Duplicates
35,789 particles
6
class (iter 36)
class (iter 37)
particles
7 - Helix Refine
90°
Resolution, loose mask
13Å
3DFSC 10.8Å
100Å
7
GSFSC Resolution: 8.64Å
Elevation
Azimuth

**Figure S6. Jurkat cell intermediate filament cryo-ET map reconstruction pipeline.** The cryo-ET map was reconstructed using cryoSPARC as outlined by the seven steps. **1** – Movie import, patch motion correction, exposure curation, and patch CTF estimation.; **2** – Manual particle picking, 2D class averages, and selection of classes; **3** – Filament tracing, inspect picks, extract particles from micrographs, 2D classification, and selection of 3 classes; **4** – Filament tracing, inspect picks, extract particles from micrographs, 2D classification, and selection of 5 classes; **5** – Helix refine; **6** – 3D classification and remove duplicate particles; **7** – Helix refine, 3DFSC, and rendering of the cryo-ET map with ChimeraX. Representative images of outputs are shown for **1** – Patch motion correction and patch CTF; **2** – Manual particle picking and three 2D class averages; **3** – Filament tracing and three 2D class averages; **4** – Five 2D class averages; **5** – Real space slices, viewing direction distribution, and FSC curves; **6** – Class flow matrix and real space slices for the five 3D classes; **7** – Real space slices, viewing direction distribution, and FSC curves.

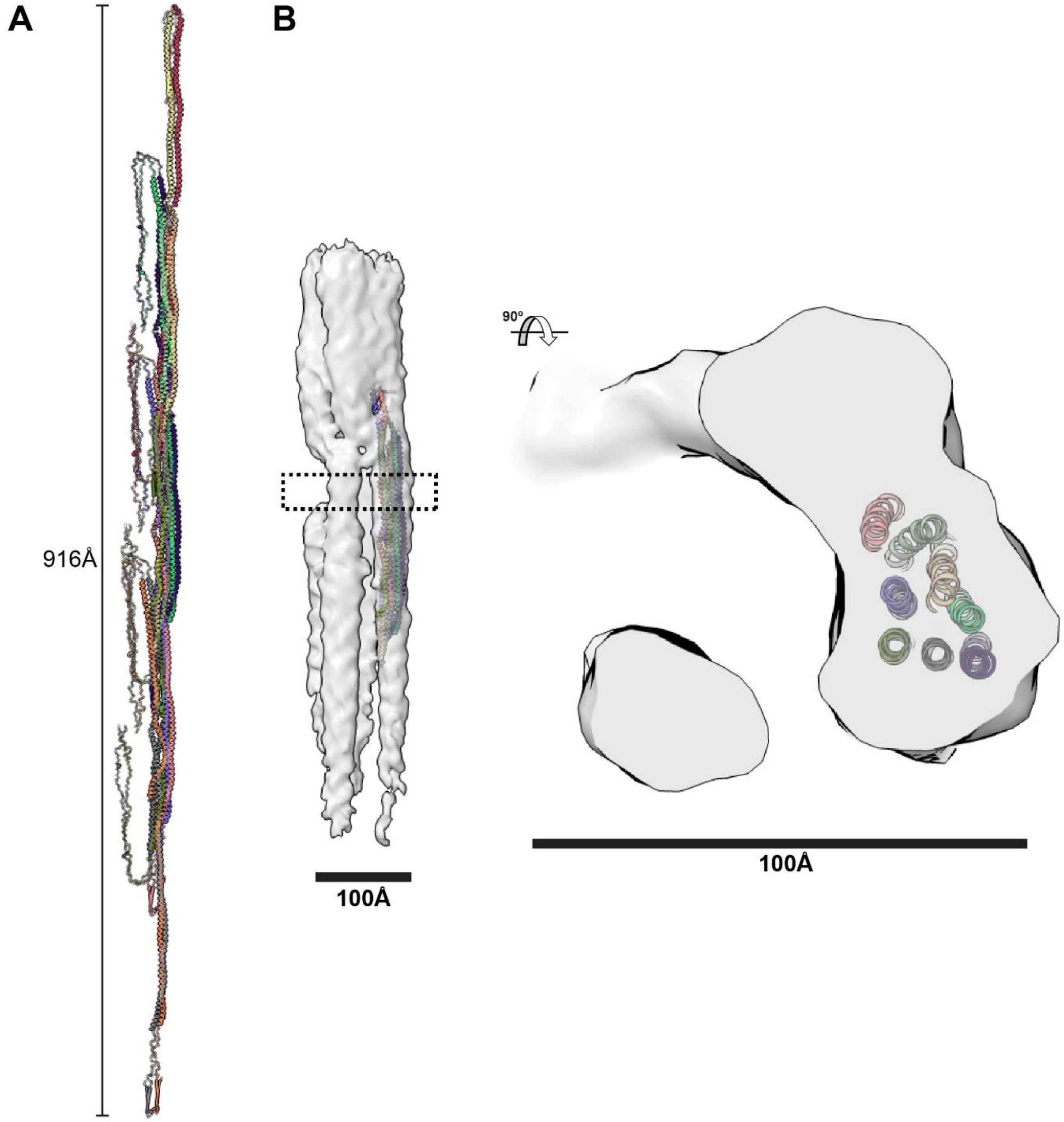


**Figure S7. The Jurkat cell intermediate filament cryo-EM map can accommodate the eight alpha helices of the vimentin protofibril. A** – Vimentin protofibril AlphaFold model reported by Eibauer et al., 2024 (PDBEV accession number PDBEV-00000212). The vimentin protofibril model is composed of three successive tetramers, creating a minimal-length protofibril with the basic repeating unit of eight chains in the center. **B** – The central eight chains of the vimentin protofibril model placed in the Jurkat cell intermediate filament cryo-EM map using ChimeraX.  The cross-section view shows the eight alpha helices of the vimentin protofibril accommodated by the cryo-EM map.  Scale bars represent 100Å.

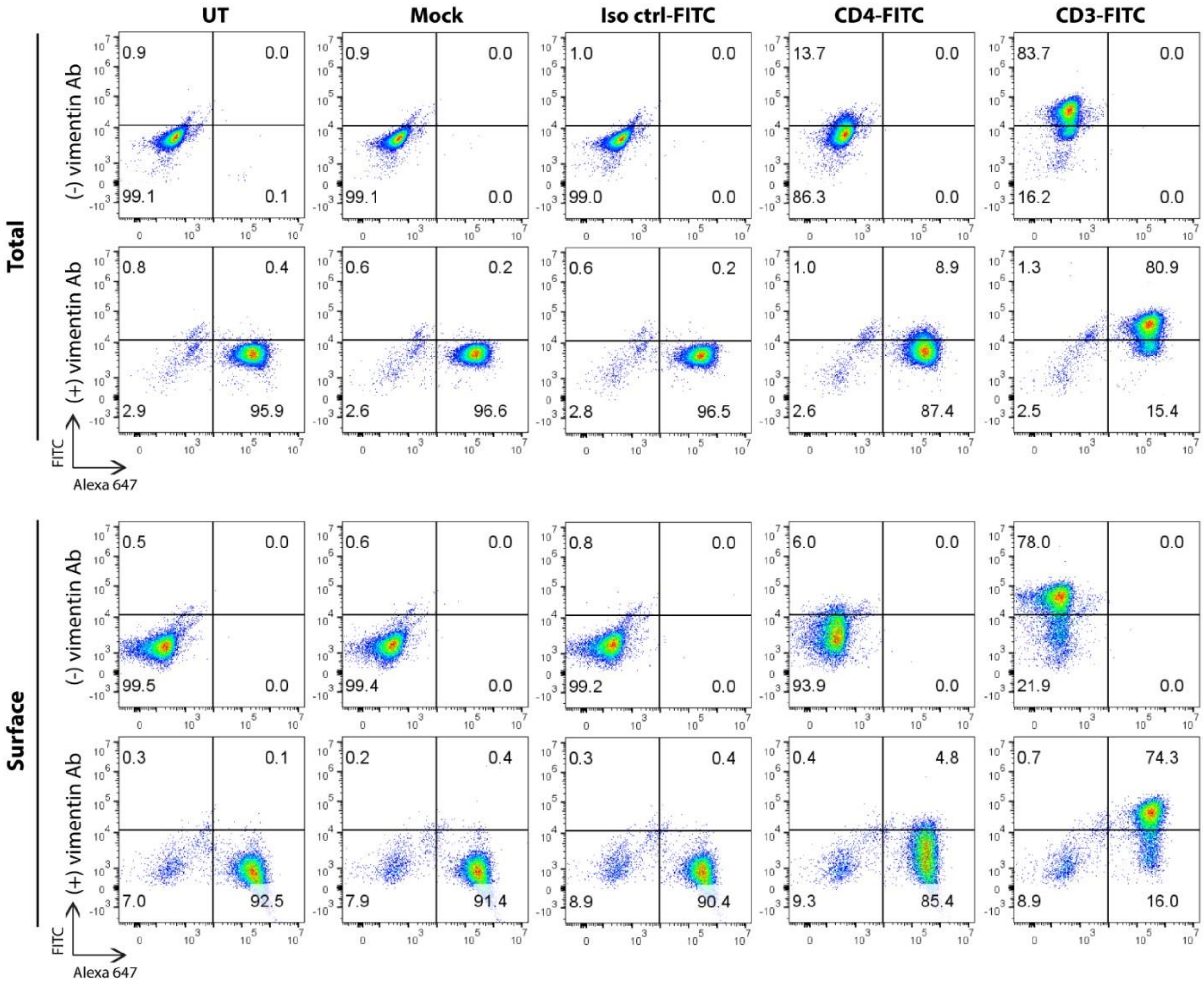


**Figure S8. Extracellular vimentin detected by flow cytometry.** Jurkat T cells preincubated with isotype control-FITC antibody (Iso ctrl-FITC), CD4-FITC antibody (CD4FITC), CD3-FITC antibody (CD3-FITC), mock treated (Mock), or left untreated (UT) at 37 ºC for 1 hr were fixed, permeabilised (Total) or left unpermeabilised (Surface), and immunostained with rabbit anti-vimentin-conjugated Alexa 647 antibody (cell signaling #9856). No antibody staining was set up as negative control for gating. Frequencies (percentage) of positive cell populations are given in each quadrant.

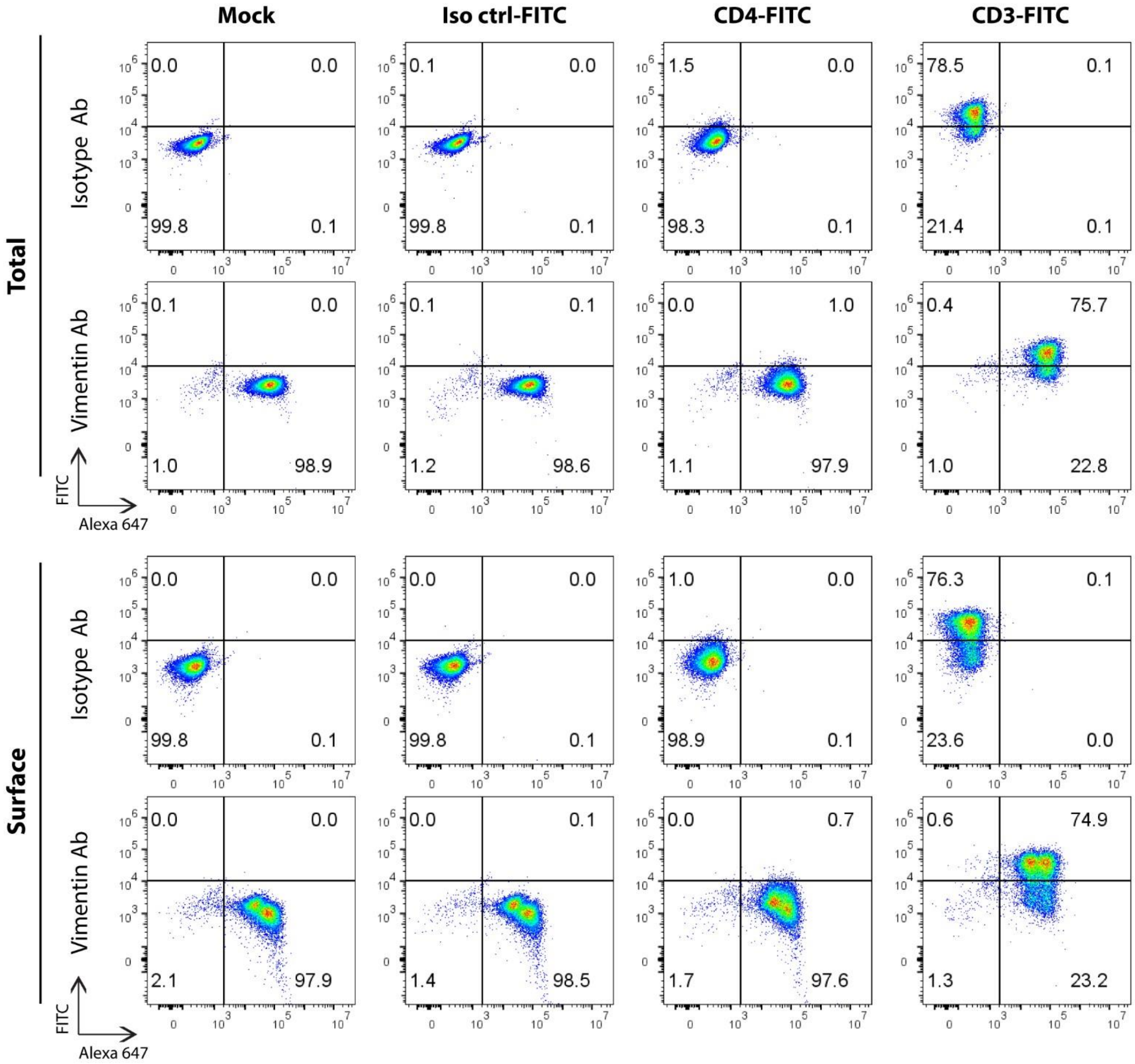


**Figure S9.** Jurkat T cells preincubated with isotype control-FITC antibody (Iso ctrl-FITC), CD4FITC antibody (CD4-FITC), CD3-FITC antibody (CD3-FITC), or mock treated (Mock), at 37 ºC for 1 hr were fixed, permeabilised (Total) or left unpermeabilised (Surface), and immunostained with rabbit anti-vimentin-conjugated Alexa 647 mAb antibody (cell signaling #9856) or with concentration-matched rabbit (DA1E) mAb Isotype ctrl-conjugated Alexa 647 (cell signaling #2985). Frequencies of positive cell populations in each quadrant are shown.

**Supplemental Table 1. Frequency of cell structures and filaments within cryo-ET montages of cryo-FIB milled Jurkat cells.**

| Grid | Lamella | Nucleus | Nuclear Pores | Cytoplasm | Mitochondria | Filaments |
|---|---|---|---|---|---|---|
| 1 (GKT) | GKT_lam1 | Yes | Yes | Yes | Yes | No |
| | GKT_lam2 | Yes | Yes | Yes | No | Yes |
| | GKT_lam3 | Yes | Yes | Yes | No | No |
| | GKT_lam5 | Yes | No | Yes | No | No |
| | GKT_lam6[A] | Yes | Yes | Yes | No | Yes |
| | GKT_lam7_2 | Yes | Yes | Yes | No | Maybe |
| 2 (VZV53G1) | VZV53G1_lam1_2 | Yes | Yes | Yes | No | Yes |
| | VZV53G1_lam2_3 | Yes | Yes | Yes | Yes | Yes |
| | VZV53G1_lam3 | Yes | No | Yes | Yes | No |
| | VZV53G1_lam4 | No | No | Yes | No | Yes |
| | VZV53G1_lam5_4 | Yes | Yes | Yes | No | No |
| | VZV53G1_lam6 | Yes | Yes | Yes | No | Yes |
| | VZV53G1_lam7 | Yes | Yes | Yes | No | No |
| 3 (VZV53G3) | VZV53G3_lamella01 | Yes | Yes | Yes | No | Yes |
| | VZV53G3_lamella02 | Yes | Yes | Yes | No | Yes |
| | VZV53G3_lamella03[A] | Yes | Yes | Yes | No | Yes |
| | VZV53G3_lamella04 | Yes | No | Yes | Yes | No |
| | VZV53G3_lamella05 | Yes | Yes | Yes | No | No |
| | VZV53G3_lamella06 | Yes | Yes | Yes | No | Yes |
| | VZV53G3_lamella07 | Yes | Yes | Yes | No | Yes |
| | VZV53G3_lamella08 | Yes | Yes | Yes | No | Yes |
| | VZV53G3_lamella09 | Yes | Yes | Yes | No | Yes |
| | VZV53G3_lamella10 | Yes | Yes | Yes | No | Yes |
| | Total (n of n) | 22 of 23 | 19 of 23 | 23 of 23 | 4 of 23 | 14 of 23 |

[A] Lamella GKT_lam6 and VZV53G3_lamella03 were used to collect tilt series for the tomograms shown in Fig. 4 and Fig. S5.

**Supplemental Table 2. Intra-bundle filament dimensions.**

| Tomogram[A] | Bundle ID | Filament Frequency | Bundle Width (nm; [SEM])[B] | Intra-bundle Filament Distance (nm; SEM)[C] |
|---|---|---|---|---|
| GKT_lam6_tilt1 | 1 | 4 | 38.39 [0.596] | 8.82 [0.163] |
| | 2 | 3 | 30.31 [0.462] | 8.49 [0.193] |
| | 3 | 2 | 22.10 [0.171] | 8.05 [0.223] |
| VZV53G3_lamella03_tilt02 | 1 | 3 | 27.58 [0.600] | 12.31 [0.389] |
| | 2 | 7 | 35.10 [0.291] | 12.56 [0.838] |

[A] Reconstructed in IMOD

[B] Measured from membrane to membrane at 3 points

[C] Measured from the filament center.

**Supplemental Table 3. Normalized RNA-seq reads from HELFs and Jurkat cells for intermediate filament genes.**

| Gene[A] | Type[B] | HELF[C] | | Jurkat[C] | | Gene[A] | Type[B] | HELF[C] | | Jurkat[C] | |
|---|---|---|---|---|---|---|---|---|---|---|---|
| | | 1 | 2 | 1 | 2 | | | 1 | 2 | 1 | 2 |
| KRT1 | I/II | 0 | 0 | 130 | 104 | KRT36 | I/II | 2 | 0 | 0 | 0 |
| KRT2 | I/II | 0 | 0 | 30 | 44 | KRT37 | I/II | 0 | 0 | 0 | 0 |
| KRT3 | I/II | 0 | 10 | 0 | 0 | KRT38 | I/II | 2 | 0 | 0 | 0 |
| KRT4 | I/II | 8 | 4 | 0 | 0 | KRT39 | I/II | 2 | 10 | 0 | 0 |
| KRT5 | I/II | 0 | 0 | 0 | 0 | KRT40 | I/II | 0 | 0 | 0 | 0 |
| KRT6A | I/II | 0 | 0 | 0 | 0 | KRT71 | I/II | 0 | 0 | 0 | 0 |
| KRT6B | I/II | 0 | 0 | 0 | 0 | KRT72 | I/II | 0 | 0 | 2 | 0 |
| KRT6C | I/II | 0 | 0 | 0 | 0 | KRT73 | I/II | 0 | 0 | 0 | 0 |
| KRT7 | I/II | 188 | 196 | 0 | 0 | KRT74 | I/II | 0 | 0 | 0 | 0 |
| KRT8 | I/II | 617 | 616 | 0 | 0 | KRT75 | I/II | 0 | 0 | 0 | 0 |
| KRT9 | I/II | 4 | 0 | 0 | 0 | KRT76 | I/II | 0 | 0 | 0 | 0 |
| KRT10 | I/II | 913 | 891 | 449 | 441 | KRT77 | I/II | 0 | 0 | 0 | 0 |
| KRT12 | I/II | 0 | 0 | 0 | 0 | KRT78 | I/II | 0 | 0 | 0 | 0 |
| KRT13 | I/II | 2 | 0 | 0 | 0 | KRT79 | I/II | 0 | 0 | 0 | 2 |
| KRT14 | I/II | 6 | 18 | 1 | 0 | KRT80 | I/II | 214 | 120 | 0 | 0 |
| KRT15 | I/II | 214 | 189 | 0 | 0 | KRT81 | I/II | 0 | 4 | 0 | 0 |
| KRT16 | I/II | 20 | 4 | 0 | 0 | KRT82 | I/II | 0 | 0 | 0 | 0 |
| KRT17 | I/II | 14 | 33 | 0 | 0 | KRT83 | I/II | 0 | 0 | 0 | 0 |
| KRT18 | I/II | 1989 | 1575 | 0 | 4 | KRT84 | I/II | 0 | 0 | 0 | 0 |
| KRT19 | I/II | 4061 | 4580 | 3 | 0 | KRT85 | I/II | 0 | 0 | 0 | 0 |
| KRT20 | I/II | 0 | 0 | 0 | 0 | VIM | III | 86069 | 81483 | 4995 | 5069 |
| KRT222 | I/II | 0 | 0 | 0 | 0 | DES | III | 77 | 43 | 0 | 0 |
| KRT23 | I/II | 2 | 1 | 0 | 0 | GFAP | III | 8 | 2 | 0 | 0 |
| KRT24 | I/II | 0 | 0 | 0 | 0 | PRPH | III | 0 | 0 | 0 | 0 |
| KRT25 | I/II | 0 | 0 | 0 | 0 | PRPH2 | III | 100 | 66 | 0 | 0 |
| KRT26 | I/II | 0 | 0 | 0 | 0 | NEFL | IV | 2 | 2 | 0 | 4 |
| KRT27 | I/II | 0 | 0 | 0 | 0 | NEFM | IV | 6 | 6 | 4 | 4 |
| KRT28 | I/II | 0 | 0 | 0 | 0 | NEFH | IV | 281 | 271 | 11 | 11 |
| KRT31 | I/II | 0 | 0 | 0 | 0 | INA | IV | 127 | 158 | 202 | 233 |
| KRT32 | I/II | 14 | 2 | 0 | 0 | LMNA | V | 19699 | 21109 | 17 | 16 |
| KRT33A | I/II | 0 | 0 | 0 | 0 | LMNB1 | V | 2110 | 1988 | 9229 | 8582 |
| KRT33B | I/II | 72 | 24 | 0 | 0 | LMNB2 | V | 5213 | 4999 | 4381 | 3973 |
| KRT34 | I/II | 527 | 487 | 0 | 0 | NES | VI | 6802 | 6475 | 9 | 8 |
| KRT35 | I/II | 0 | 0 | 0 | 0 | | | | | | |

[A] Gene name symbol.

[B] Intermediate filament type.

[C] Normalized RNA-seq reads from duplicate samples.

**Supplemental Table 4. Cryo-ET data collection parameters**

| Parameter | Titan Krios (FEI; TEM1) |
|---|---|
| Voltage (kV) | 300 |
| Magnification | 26,000 |
| Gatan Detector (pixels) | K3 (5760 4092) |
| Energy filter slit width (eV) | 20 |
| Pixel size (Å) | 3.465 |
| Defocus range (μm)[A] | -2 to -4 |
| Exposure time (s) | 2.49667 |
| Frames | 10 |
| Dose Rate $e^-$/pixel/s | 8.44102 |
| Electron dose ($e^-/Å^2$/frame) | 0.1755 |
| Electron exposure rate ($e^-/Å^2/s$) | 0.70305 |
| Tilt angle (°±) | 0-60 |
| Tilt step (°)[B] | 2 |
| Total electron exposure per frame ($e^-/Å^2$) | 1.755 |
| Total electron exposure per tilt series ($e^-/Å^2$) | 107.073 |

[A] Target defocus

[B] Dose symmetric
