## Supplemental Movie Legends for "Extracellular filaments revealed by affinity capture cryo-electron tomography"

**Supplementary Movies 1 to 3. Aligned tilt series of extracellular intermediate filaments generated by cryo-FIB/cryo-ET.** These movies show aligned tilt series generated by cryo-FIB/cryo-ET that reveal extracellular filaments at the periphery of Jurkat cells. Dense networks of filaments in the extracellular space, bundles of filaments emerging from the plasma membrane, and the intracellular and extracellular environments are labelled.

**Supplementary Movie 4. Extracellular intermediate filaments revealed by cryo-FIB/ET.** The movie shows a reconstructed cryo-tomogram of extracellular intermediate filaments at the periphery of a Jurkat cell. A single slice of the tomogram is show in **Fig. 2C**. The scale bar (black) in the bottom right-hand corner represents 100nm.

**Supplementary Movie 5. AI-based segmentation of extracellular intermediate filaments.** The movie shows reconstructed cryo-tomograms for two representative Jurkat cells and AI-segmentation of the extracellular intermediate filaments (green), plasma membrane (brown), intracellular vesicles (yellow), and the nuclear envelope (Red). Single slices of the cryo-tomograms are shown in **Fig. 5B and 5D**. The scale bar (white) in the bottom right-hand corner represents 100nm.
